# Multi-histone ChIP-Seq Analysis with DecoDen

**DOI:** 10.1101/2022.10.18.512665

**Authors:** Tanmayee Narendra, Giovanni Visonà, Crhistian de Jesus Cardona, Gabriele Schweikert

## Abstract

Epigenetic mechanisms coordinate packaging, accessibility and read-out of the DNA sequence within the chromatin context. They significantly contribute to the regulation of gene expression. Thus, they play fundamental roles during differentiation on the one hand and maintenance and propagation of cell identity on the other. Epigenetic malfunctioning is associated with a large range of diseases, from neurodevelopmental disorders to cancer progression. In humans, hundreds of known epigenetic factors and complexes are involved in establishing covalent modifications on the DNA sequence itself and on associated histone proteins. Within the cellular context, the resulting combinatorial epigenomic patterns are neither established nor interpreted independently of each other and therefore exhibit high correlations in a region-specific manner. Post-translational modifications of histone proteins can be analysed using Chromatin Immunoprecipitation followed by sequencing (ChIP-Seq). Often, several assays for a number of different histone modifications are performed as part of the same experimental design. These measurements are, however, confounded by shared biases including chromatin accessibility and mappability. Existing computational methods analyse each histone modification separately. We introduce DecoDen, a new approach that leverages replicates and multi-histone ChIP-Seq experiments for a fixed cell type to learn and remove shared biases. DecoDen (Deconvolve and Denoise) consists of two major steps: We use non-negative matrix factorisation (NMF) to learn a joint cell-type specific background signal. Half-sibling regression (HSR) is then used to correct for these biases in the histone modification signals. We demonstrate that DecoDen is a robust and interpretable method that enables the unbiased discovery of subtle peaks, which are particularly important in an individual-specific context.

## 1 Introduction

The combination of Chromatin Immunopreciptation assays with sequencing (ChIP-seq) is a powerful tool to identify transcription factor binding sites and enrichment of histone modifications [7].

Experimental protocols for histone ChIP-Seq broadly follow these steps as depicted in Figure 1 A [12]: Chromatin is crosslinked by foramldehyde to stabilize the interactions between DNA and associated proteins. This step is sometimes omitted and chromatin is used in its native state. Chromatin is then sheared into managable fragments through sonication. An antibody specific to the protein of interest is chosen to immunoprecipitate the DNA-protein complex. Next, the crosslinks are reversed. After selection of fragments in an optimal size range of 200-300 bp, the resulting DNA fragments are sequenced either from one and or both producing either single-end or paired end reads of a given read length, i.e. 50-100 bp.

**Fig. 1.**
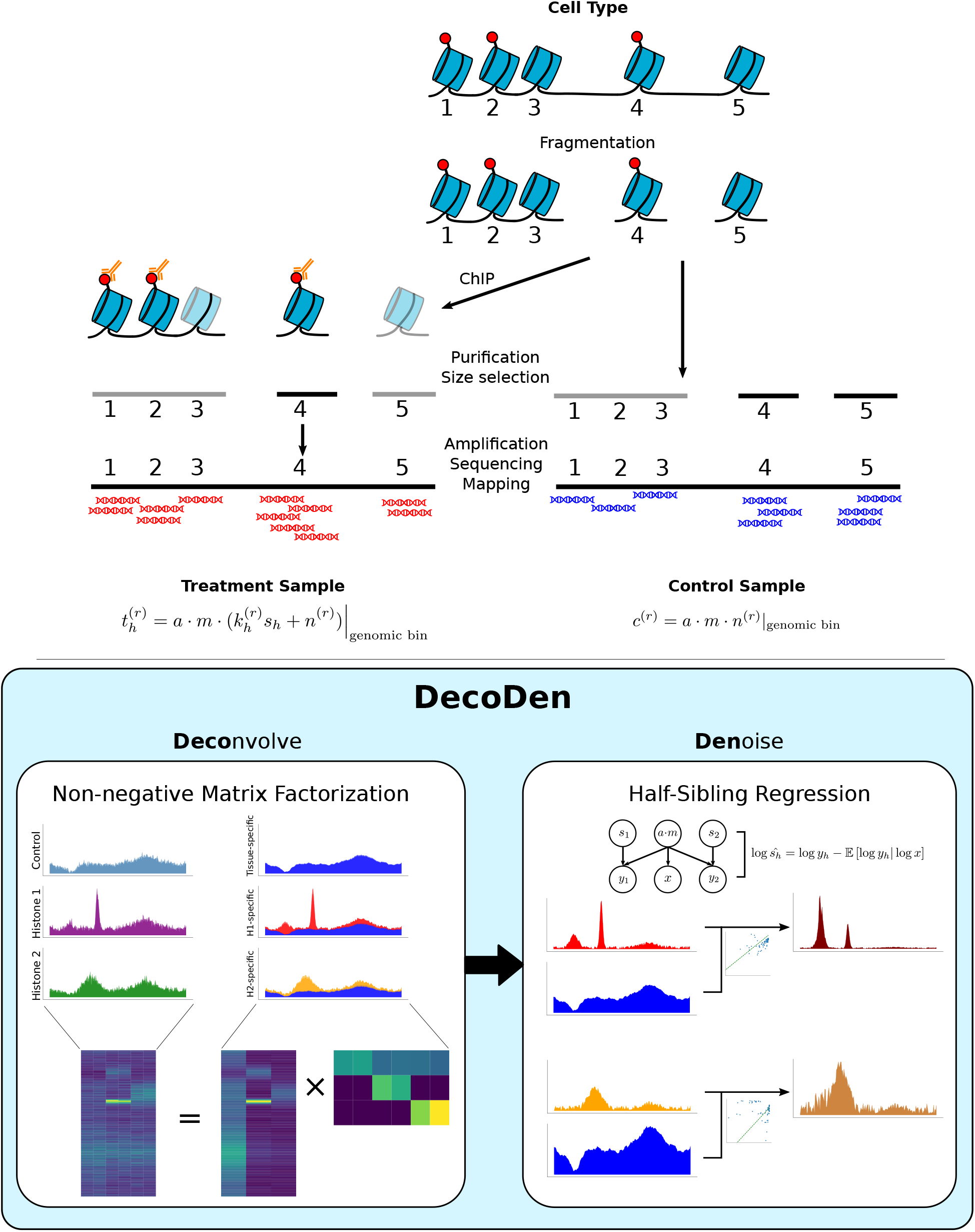
Schematic of DecoDen. A. Experimental ChIP-Seq procedure. B. Computational steps of DecoDen.

There are several sources of bias in a ChIP-Seq experiment. First, as open chromatin regions are more easily sheared than other regions cell-type specific chromatin accessibility biases arise during genome sonication, where open regions yield more protein-DNA complexes. Second, the antibody used for immuno-precipitation has an inherently variable specificity, which can lead to pileups of off-target reads in the sequencing process which obfuscate the enriched regions [3]. There are other sources of variability such as sequence-dependent PCR duplication, and copy number variation, which are specific to the cell-type under study and independent of the protein of interest. In order to capture these cell-type specific biases, most ChiP experiments are accompanied by a control experiment where input DNA prior to immunopreciptation is assayed [12], see Figure 1A) [12].

The analysis of ChIP-seq data is a complex multistep process that starts from sequenced reads and ends with the prediction of regions with specific histone modifications, transcription factor (TF) binding sites, or chromatin marks. *Peak calling* involves identifying regions that are significantly enriched for the protein of intresst in the ChIP-ed sample relative to the control. This is an imbalanced data problem, since enriched regions generally span less than 5% of the genome. In addition, the fraction of reads in enriched regions (FRiP) is less than 10% in the ChIP-ed sample and even less in the control. This makes the detection of subtle variations in signal difficult to detect[18].

A plethora of bioinformatics software packages have been developed for the task of peak calling, such as diffReps [17], SICER [19], and PePr [20]. The most cited among them, however, is MACS (Model-based Analysis of ChIP-Seq) [21]. MACS is usually the default tool for ChIP-Seq analysis, and large-scale studies such as the ENCODE project [4] and the Roadmap Epigenomic Project [9] use it in their data processing pipelines.

MACS identifies enriched regions in the following steps. Consider two samples - the treatment sample with reads from ChIP enrichment and the control sample with reads from the control experiment. In the first step of MACS, duplicate reads are removed, fragment length is estimated and reads are extended up to the fragment length. This is done for both treatment and control samples. The next step is to compute the local background distribution from the control sample against which the treatment sample will be compared. To compute the background at each genomic base, MACS takes the maximum value of the read counts at that base, and that at windows of 1 kB, 10 kB and the genome background. Since the genome background is defined as 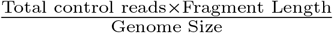, it is ensured that all bases have a non-zero value for the background. Next, the treatment and the control samples are normalized to the same total sequencing depth. To compute enrichment of treatment over background, a Poisson significance test is done at every base. This results in a genome-wide signal of p-values of enrichment. Contiguous regions with an enrichment greater than a specific cut-off are specified as peaks [1, 21].

This procedure has some drawbacks. The mean parameter of the local background Poisson distribution is estimated from a single control measurement. The Poisson distribution is asymmetric, and this asymmetry is particularly pronounced for lower values of the mean. Hence, using a single sample to estimate the mean of the background at a base leads to underestimating the value of the true mean more often (Supplementary Figure 1). Another issue arises from insufficient sequencing depth. Many ChIP protocols (for example, the Roadmap Epigenomic project [9]) use comparable library sizes for both the ChIP and control samples, even though a control sample would require a higher library size, since it aims to profile experimental bias for the entire genome. When the control sample has insufficient sequencing depth, it results in insufficient coverage of the genome and the background distribution is underestimated. With MACS, this increases the likelihood of using the genome background as an approximation of the true signal. As a consequence, the background is underestimated in regions with high accessibility and mappability and underestimated in regions where accessibility and mappability are low. As a consequence, on average, histone marks associated with closed chromatin have lower maximum significance than those associated with open chromatin (Supplementary Figure 3). When all histone marks are sequenced to the same library size, this induces a systematic bias against histone marks associated with closed chromatin. Histone modifications interact in complex ways to regulate gene expression. Hence, it is often the case that more than one histone modification is assayed in a given cell-type. When the goal of an experiment is to understand the interplay of histone modifications in a specific cell type, it becomes important to correctly model the biases introduced by chromatin accessibility, PCR amplification and mappability. Without this step, it is difficult to distinguish between experimental noise and true signal. While technical replicates are used in almost every experimental design, peak calling tools such as MACS do not leverage them to separate experimental noise from the true signal.

We introduce DecoDen (Deconvolve and Denoise), a new method to remove cell-type-specific bias for samples of different histone modifications derived from the same cell type samples. This method leverages different assays and jointly analyses data to remove noise and biases common to the cell type. We demonstrate that DecoDen reduces spurious peaks by sharing information across samples. DecoDen is available as an open-source tool at https://github.com/ntanmayee/DecoDen.

## 2 Results

### 2.1 The DecoDen Model

#### Data Preprocessing

Similar to MACS, we filter duplicate reads and estimate fragment length. Reads are extended in the 5’ to 3’ direction for ChIP reads, and in both directions for input reads, up to the estimated fragment length. For faster computation, average coverage is computed for fixed-size bins. In our experience, bin sizes varying from 10 to 200 are reasonable. Anything higher than this can fail to distinguish between signal and noise. For analysis with real experimental data, the *Low Mappability* regions from the ENCODE blacklist [2] were removed. The resulting data is given as input to subsequent steps.

DecoDen consists of two steps. The first step uses non-negative matrix factorization to separate the cell-type specific signal from the ChIP signals specific to the different histone modifications. Next, we use Half-Sibling Regression to isolate the true (hidden) enrichment pattern by minimizing the bias introduced by accessibility and mappability. A conceptual overview of DecoDen can be seen in Figure 1.

Let us assume that the observed data 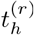 from an experimental assay for the *h*th histone modification and *r*th replicate is a vector of dimension *G*, the length of the genome. Given that antibodies are not 100% specific, fragments that pass the Immuno-precipitation step will be a mixture of target fragments and background. 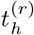 can be written as

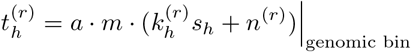

where *a* captures local cell-type specific biases including chromatin accessibility, and *m* captures biases which depend on the depend on the DNA sequence alone, including mappability and amplification bias. *s*_*h*_ is the true underlying enrichment for the *h*th histone modification, 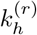 is the mixing parameter and *n*^(*r*)^ is the unspecific noise. Similarly, the input sample can be written as

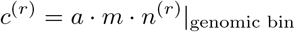

#### Non-negative Matrix Factorisation

Non-negative matrix factorisation (NMF) is a general methodology to decompose a given matrix V with dimensionality *i* × *j* into two matrices *W* and *H* with dimensionality *i* × *l* and *l* × *j* such that:

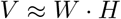

where · denotes the scalar product.

The motivation behind using NMF is to separate histone-specific signals from unspecific additive noise. In particular, all ChIP and input measurements are analysed together to deconvolve them into a consolidated signal for each histone modification *a m s*_*h*_ and an unspecific noise component *a m n* that captures cell-type -specific and -unspecific biases. Since this step explicitly uses replicates to share information across samples, experimental noise among samples is minimised. Unlike MACS, NMF looks for correlations on a global rather than a local scale, thereby inducing information sharing among genomic loci. In DecoDen, the *V*_*i×j*_ matrix consists of a concatenation of treatment and input samples, where 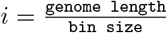 and *j* is the total number of samples. The *l* parameter is chosen based on the number of different kinds of measurements that are present. (For example, *l* = 3 when there are measurements for H3K4me3, H3K27me3 and control.) We thus learn specific signals for each histone modification *y*_*h*_ = *a* · *m* · *s*_*h*_, and an additional control signal *x*, containing valuable information about shared biases *a · m*. These biases affect both control and ChIP samples, and *x* is thus used in subsequent steps to de-bias the assay-specific signals.

Non-negative matrix factorisation is NP-hard in general, and convergence heuristics are dependent on initialisation [22]. To ensure that the deconvolution step captures the desired information, we split the matrix factorisation into three steps. Firstly, we factorise only the signals for the control replicates, which results in learning a cell type-specific background and the mixing matrix parameters for the input control samples. Subsequently, we fix this background signal *x* and extract the mixing coefficients 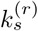 for the treatment sample that minimize the loss function, to “explain” as much of the treatment signal as possible using only the control signal. Finally, we subtract from the treatment signal this background component and learn the remaining mixing coefficients jointly with the histone-specific signals *y*_*h*_, to explain the remaining information. The optimisation is carried out by minimising the KL Divergence as loss function.

#### Half-Sibling Regression

Half-sibling regression [16] is a method inspired by causal inference to remove the effect of confounders. Chromatin accessibility is a known confounder in estimating the true enrichment of a histone modification. For instance, the highest values for ChIP tracks are higher for histone marks associated with open chromatin when compared to those associated with closed chromatin (See Supplementary Figure 3). Mappability is a measure of the number of unique genome positions that a *k*-mer can be aligned against. It is sequence-specific, and its variation confounds comparison of different genomic regions. In our approach, we model a cell-type-specific confounder as a combination of accessibility and mappability. (Refer to the directed graph in Figure 1) For a given cell type, this confounder is fixed, and affects both the control sample and the histone specific signals. In addition, we make the simplifying assumption that the true underlying histone modification signal is independent of the cell-type specific signal or *s*_*h*_ ╨*x*. This allows us to use half-sibling regression to remove the effect of confounding factors between the ChIP signal and the cell-type specific background signal. The true histone modification signal *s*_*h*_ is estimated as

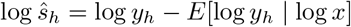

where *y*_*h*_ is the histone-specific signal estimated from NMF and *x* is the cell-type specific background signal estimated from the previous NMF step.

The application of HSR to logarithm-transformed signals using a linear regression model allows us to filter multiplicative noise with better numerical stability. Since subtraction in the log scale translates to division in the linear scale, this is similar to computing fold-change. The estimated signal *ŝ*_i_ is recovered by exponentiating the estimate from half-sibling regression.

Since the HSR step is equivalent to dividing by a confounding factor, certain genomic loci may present extreme values due to a background signal that is essentially 0. To avoid this pitfall, we bounded the divisor to a minimum value of 0.5 (i.e. capping the predictions of the fit in log scale to *log*(0.5)), essentially ensuring that the signal is at most amplified by a factor of two.

#### Identifying enriched regions

We use the estimated fold-change from the previous Half-Sibling Regression step and a fixed threshold to identify regions of enrichment. Regions that are close together are merged, and extremely small enriched regions are eliminated.

### 2.2 DecoDen recovers subtle peaks by sharing information across samples and experimental conditions

DecoDen jointly analyses different experimental conditions and replicates. While it is possible to specify different replicates with MACS, it internally merges all treatment and control replicates respectively and loses the ability to distinguish between signal and technical noise. Figure 2A shows results from peak calling with DecoDen and MACS for a dataset simulated with ChIPulate [6]. (Details about the simulation are in *Methods*). For DecoDen, intermediate results from non-negative matrix factorisation and half-sibling regression are shown. For MACS, the enrichment is provided as –log_10_ (p-value) and is tiled into the same bin width as DecoDen. While it is evident that MACS and DecoDen provide similar results, DecoDen highlights sharper peaks and is better able to distinguish between signal and noise. The rest of the figure shows specific examples where there is a mismatch between DecoDen and MACS. Figure 2B is an example where merging replicates results in MACS identifying false peaks (left and right extremes) while missing a true peak (center). Figures C is an example where MACS incorrectly estimates a lower significance for a true peak and a higher significance for a region without peaks. Figure D and E are more examples showing how MACS over-estimates enrichment.

**Fig. 2.**
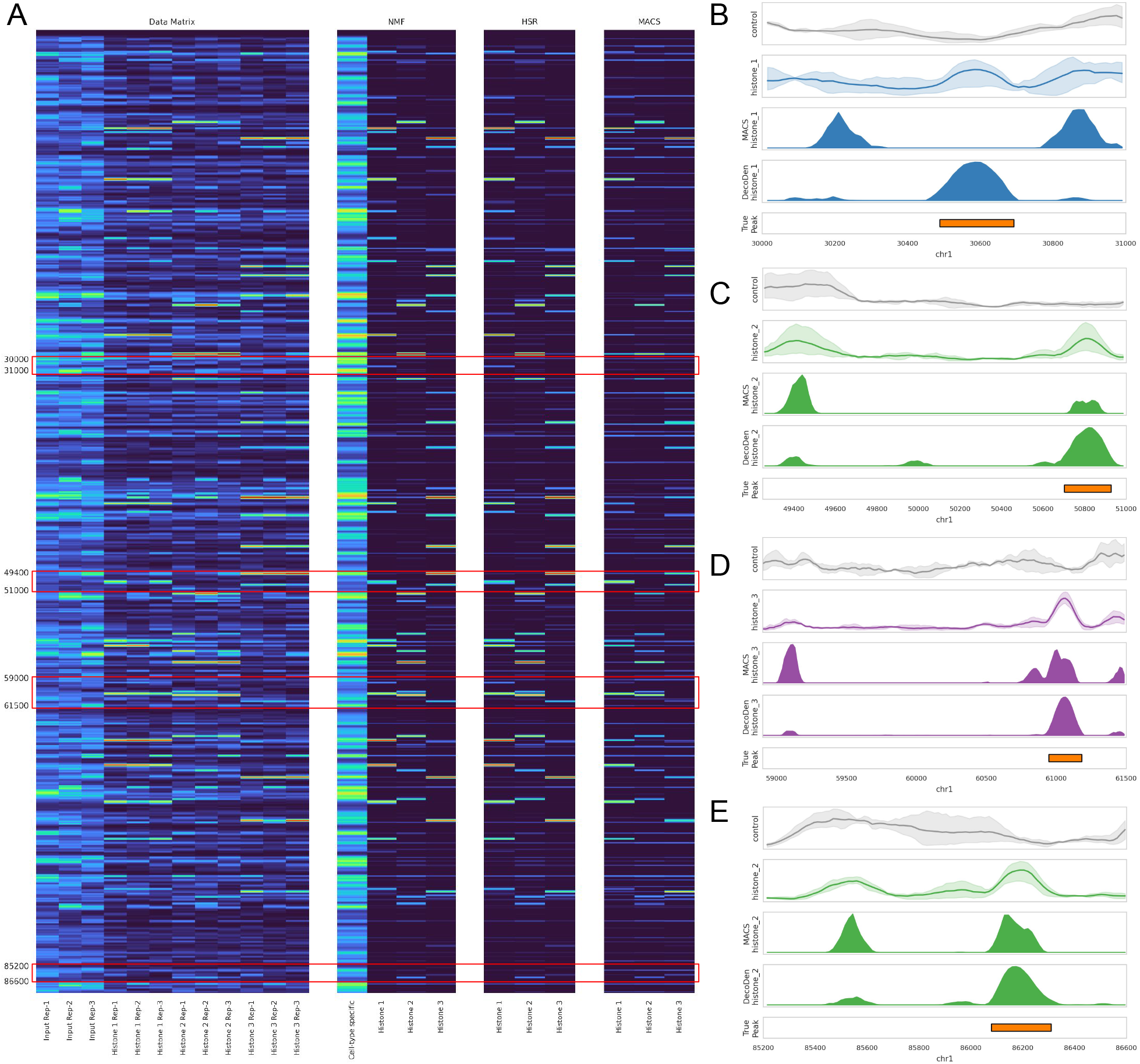
Comparison of DecoDen and MACS for simulated data: A. Heatmap of simulated regions using the turbo colour map to highlight disparities. B-E: examples of regions of mismatch. The first two rows show mean coverage and variation across replicates.

### 2.3 DecoDen removes mappability and sonication biases

Recall that the learning scheme is set-up in such a way as to explain most of the genomic bins in the ChIP samples with the input sample. (See examples of the mixing matrix in Supplementary Figure 2.) This is motivated by the fact that enriched regions constitute about 5-10% of the genome and a majority of the reads constitute the background signal. Hence, the cell-type specific background signal *x* in the *W* matrix of NMF contains information common to all replicates and conditions. We hypothesize that this captures cell-type specific information about mappability, sonication and fragmentation. Figure 3 demonstrates how the shape of the H3K4me3 enrichment is conserved while eliminating cell-type specific bias. This figure uses data where DecoDen was run on deeply sequenced data for A549 Lung Carcinoma cell line (E114) from [8] in order to avoid biases that arise from insufficient sequencing depth. To highlight the utility of DecoDen in scenarios where there is a larger number of experimental conditions and replicates available, data for H1 BMP4 Derived Trophoblast Cultured Cells (E005) from the Roadmap Consortium was used. An example of the reconstruction can be seen in Figure 4 where the signal spike common to all samples (around 44,790 Kb) is captured by the cell-type specific background signal.

**Fig. 3.**
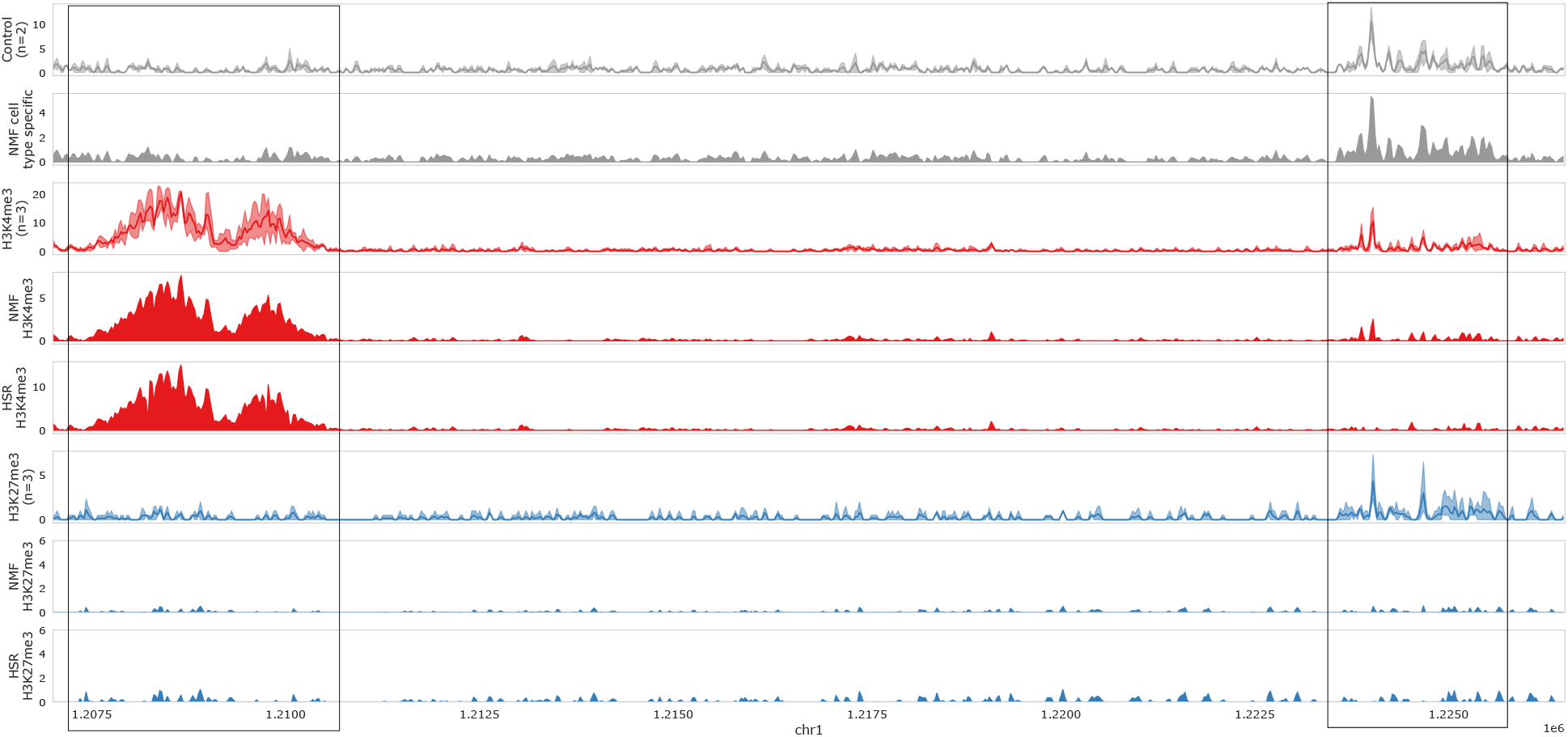
Reconstruction using DecoDen for deeply sequenced A549 Lung Carcinoma cell line (E114): While the NMF step reduces noise among replicates, the HSR step eliminates cell-type-specific bias as seen in the region around 1225 Kb. The shape of the peak is preserved as seen in the H3K4me3 peak on the left side.

**Fig. 4.**
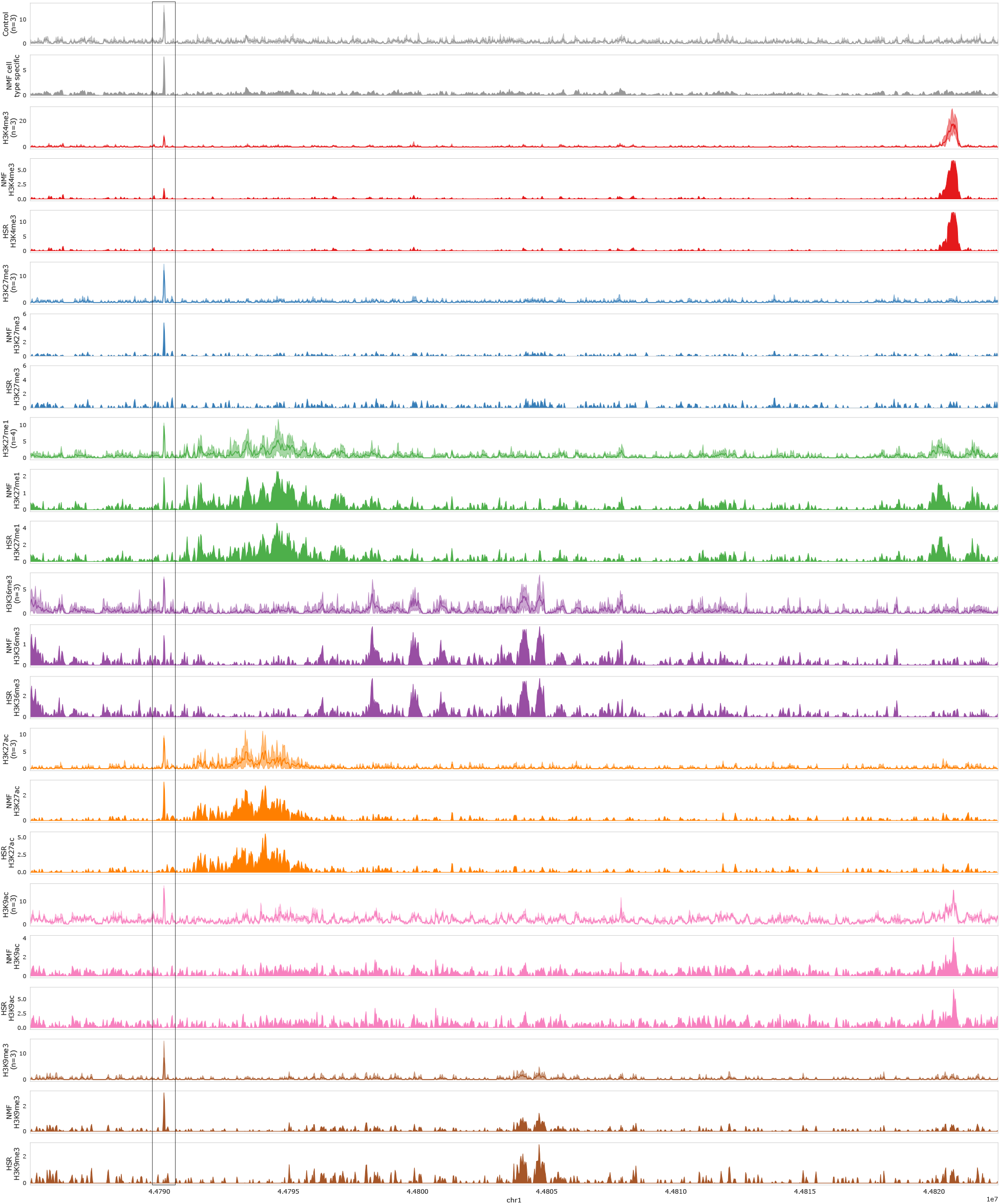
Reconstruction using DecoDen for H1 BMP4 Derived Trophoblast Cultured Cells (E005) from the Roadmap Consortium: The first row in each colour shows coverage of replicates after the pre-processing step. The second and third row show the signal after the NMF and HSR steps, respectively. The highlighted spike is identified as a cell-type specific signal in the NMF step and is removed from all histone marks after HSR.

### 2.4 DecoDen reduces correlation between Input Control and ChIP measurements

Different histone modifications require different sequencing depths to reach saturation [8]. Histone modifications associated with gene repression are often correlated with regions of low chromatin accessibility, and reach saturation at higher sequencing depths. This confounds estimation of enrichment. When data from activating and repressing histone marks are sampled to the same sequencing depth, this introduces a bias where peaks for activating marks have a higher statistical significance than those associated with repression (Figure 3 in Supplementary Material). When different histone modifications are profiled for a fixed cell type, it is important to minimize differences due to chromatin accessibility. These differences are seen clearly in Figure 5, which shows ChIP data for transverse colon tissue for individual 4 of the ENTEx dataset [15]. While all histone modifications show correlation with the control sample, those associated with gene activation such as H3K4me3 and H3K27ac have a higher number of genomic positions with a high treatment coverage and low control coverage. The genomic bins on the diagonal have comparable coverage for both treatment and control, indicating that the high coverage arises from cell-type specific biases that allows a greater propensity for sequencing. After DecoDen, the genomic bins that were previously on the diagonal are designated as cell-type specific rather than assay-specifc enriched regions.

**Fig. 5.**
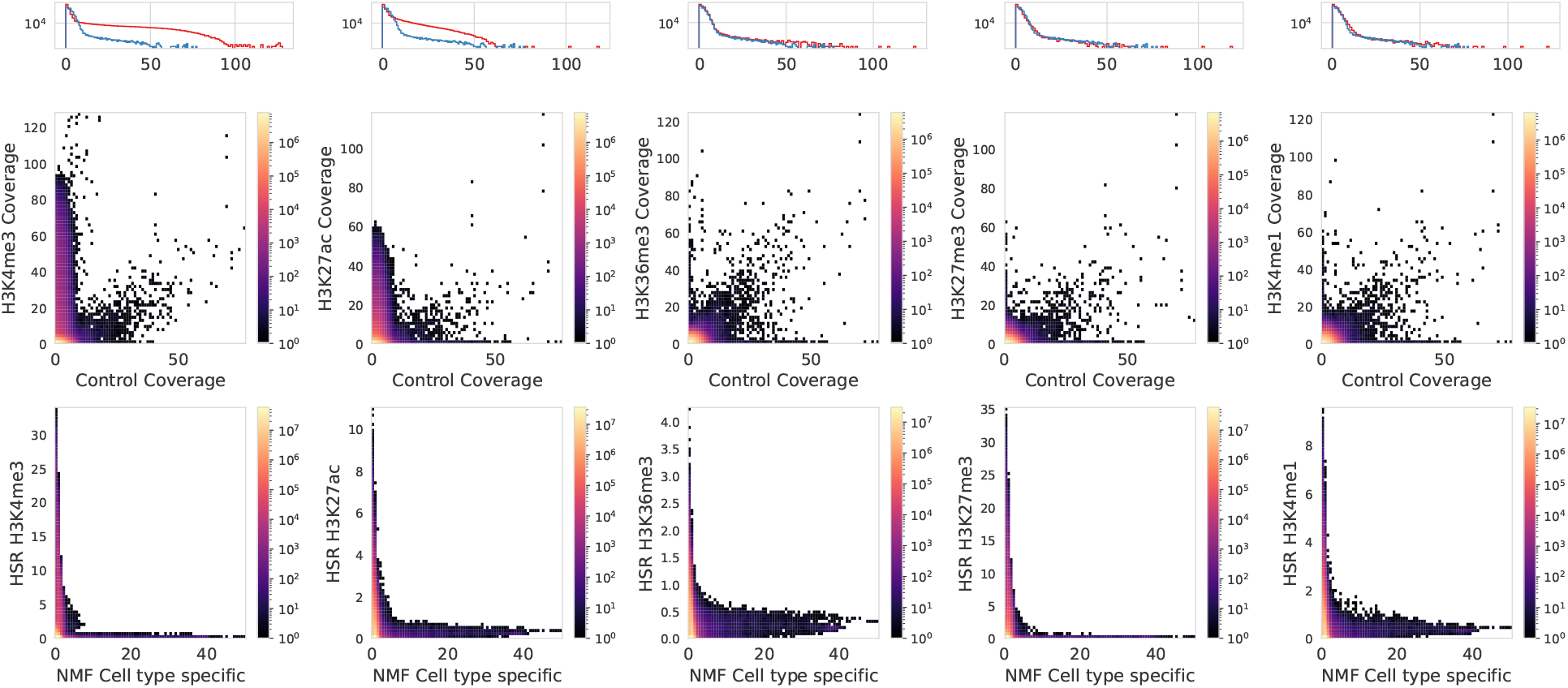
DecoDen removes correlation between control and treatment samples: Top row: histogram of coverage, red represents the histone modification and blue the input control. Second row: 2d histogram between control and treatment coverage. All samples have genomic regions along the diagonal. The bottom row shows the corresponding histogram plot after DecoDen. Data from the transverse colon tissue from Individual 4 (51F) of the ENCODE ENTEx dataset is shown.

### 2.5 Identifying individual specific peaks

When the goal is to identify histone modification patterns that are specific to an individual, it is important to minimize amplification, mappability and cell-type specific biases to better characterize smaller peaks. Consider the example of ChIP experiments for transverse colon for individuals 3 and 4 in the ENTEx consortium [4, 10, 15]. For gene activating histone marks that are more likely to reach saturation at lower sequencing depths (H3K4me3 and H3K27ac in this case), the correlation between individuals is only slightly smaller than the correlation between samples of a particular individual (See Table 1 in Supplementary Material for details). Supplementary Material Figure 4 is an example where MACS overestimates the significance for some regions, thereby distorting the data for downstream bioinformatic tasks.

## 3 Materials and Methods

### 3.1 Data

#### Simulated Data

Since true enrichment regions for experimental data are unknown, synthetic data using the ChIP-Seq simulation tool Chipulate[6] was generated. Chipulate generates ChIP and treatment reads for predefined peak regions. Each enriched region has three additional parameters to specify extraction efficiency `p_ext`, PCR amplification efficiency `p_amp` and binding energy `energy_A`. Further details regarding the simulation procedure are provided in the Supplementary Material.

#### Experimental Data

Deeply sequenced ChIP-Seq data was taken from [8]. This paper used the A549 Lung Carcinoma cell line, which corresponds to E114 in the Roadmap Consortium Dataset. We used the data for histone modifications H3K4me3 and H3K27me3 (three technical replicates each) and the whole cell extract (WCE) input sample (two replicates). SAMtools [5] was used to align the data to the hg19 genome. Picard was used for quality control of the aligned reads. In particular, reads failing vendor quality checks, those which are PCR duplicates and unmapped reads were removed. Additionally, we used data for H1 BMP4 Derived Trophoblast Cultured Cells (E005) from the Roadmap Consortium. The following histone modifications were used - H3K9me3, H3K27me3, H3K4me3, H3K4me1, H3K36me3, H3K27ac and H3K9ac in addition to WCE input. Since DecoDen leverages multiple replicates, the unconsolidated data was used. As E005 is a cell line, the data consisted exclusively of technical replicates. The data set is provided as `tagAlign` files which have been aligned to the hg19 genome. To characterize individual specific differences, data for transverse colon tissue for histone modifications H3K4me3, H3K36me3, H3K4me1, H3K27ac and H3K27me3 from the ENCODE ENTEx Project [4, 10, 15] was used. Data is provided as `bam` files aligned to the GRCh38 genome. Accession numbers are provided in Section 3 in Supplementary Material.

DecoDen internally uses BEDtools [14] to compute coverage for each sample. BEDOPS[11] is used to compute the mean coverage of each track at each genomic bin. All NMF experiments are carried out using the Python package *scikit-learn* [13]. MACS[21] commands `filterdup`, `predictd` and `pileup` are used to filter reads, predict fragment length and extend reads respectively. MACS `bdgcmp` was used to compute enrichment in – log_10_ p-value with additional parameter `-m qpois` to apply Benjamini-Hochberg correction.

### 3.2 Training and Validation

In the case of experimental datasets, to evaluate the components of the mixing matrix that describe the proportions of each signal that compose the measured replicates, we select 400,000 genomic bins (bin-width of 25 bp) as training set from regions with background coverage ≥ 1. We employed L2 regularisation on both the signal matrix and the mixing matrix; to determine the weights of the regularisation term in the loss function, we selected 10,000 additional bins to use as validation set to choose a sufficiently high regularisation that would not adversely impact the reconstruction of the signal. The final configuration of the algorithm used a weight of 0.01 for the L2 regularisation of the signal matrix and 0.001 for the regularisation of the mixing matrix. For simulated data, the same regularisation parameters were used. Since the simulated genomic region is much smaller than experimental data, the number of genomic bins for training was 5000 with a bin-width of 15 bp.

## 4 Discussion

In this paper, we introduce DecoDen (Deconvolve and Denoise), a new method to leverage replicates and different experimental assays in ChIP-Seq experiments to reduce systematic noise and cell-type-specific biases. DecoDen consists of two major steps: non-negative matrix factorization (NMF) to deconvolve the histone-specific signal from the background, and half-sibling regression (HSR) to correct the sources of bias shared between the components.

NMF is used with a custom initialization scheme to optimise the estimation of the cell-type-specific signal and histone-modification-specific signals. The mixing matrix from the NMF step is biologically interpretable, and indicates how much of the ChIP sample is histone-modification-specific, and how much is cell type specific. Since genome positions are treated together and not independently, the analysis is at a global rather than local scale. Further, all samples are analysed jointly instead of separately. This combined analysis helps in sharing information among samples and genomic positions. The HSR step, based on a causal graph of the data-generating process, removes the effect of cell-type-specific confounders such as accessibility and mappability. This procedure makes it possible to jointly analyse histone modifications requiring different saturation depths. We use simulated and experimental data to demonstrate how DecoDen identifies subtle peaks without distorting the data. This is especially important in an individual specific context.

An important caveat is that DecoDen requires a minimum of two replicates for each histone modification to converge to a non-trivial solution. Increasing the number of replicates and/or the diversity of histone modifications increases the statistical power, and hence should improve results. One possible avenue for future work is to explore how samples from other cell types can be used to potentially separate out sequencespecific biases from cell-type-specific biases, including chromatin accessibility and sonication bias.

## Acknowledgements

The authors thank the International Max Planck Research School for Intelligent Systems (IMPRS-IS) for supporting Tanmayee Narendra and Crhistian de Jesus Cardona.

