## Supplement for "Multi-histone ChIP-Seq Analysis with DecoDen"

#### 1 Simulation Procedure

We have generated synthetic data using the ChIP-Seq simulation tool Chipulate. Chipulate generates ChIP and treatment reads for predefined peak regions. Each enriched region has three additional parameters to specify extraction efficiency `p_ext`, PCR amplification efficiency `p_amp` and binding energy `energy_A`. To generate input replicates, a contiguous region of about 100 kB was defined with randomly sampled parameters for `p_ext` and `p_amp`. To generate treatment replicates, each histone modification was assigned randomly sampled regions of enrichment. The same `p_ext` and `p_amp` parameters from the input were used. Binding energy was randomly sampled from a predefined range. Three histone modifications were simulated with four replicates each. To mimic technical noise, the parameters for each replicate was perturbed by adding Gaussian noise. Chipulate generates reads only inside peak regions. To mimic realistic FRiP ratios, additional randomly sampled reads from input were added to the treatment replicates. MACS `callpeak` was run with default parameters with `--nomodel` flag.

#### 2 Supplementary Figures

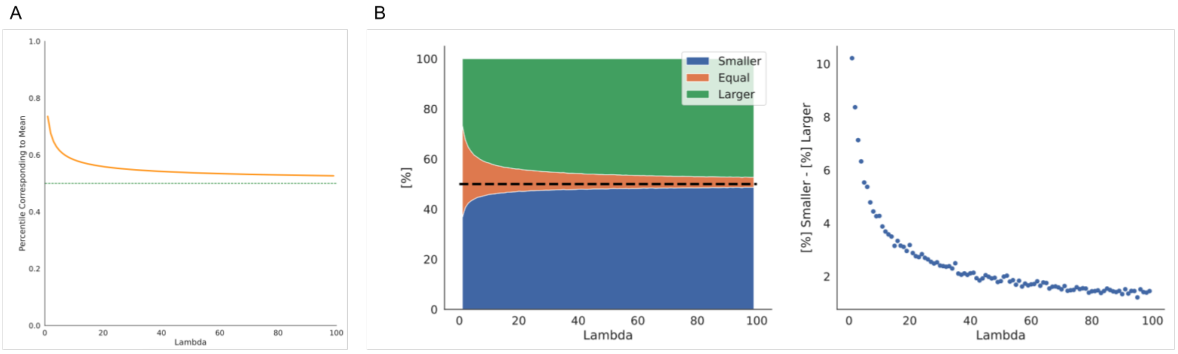


Figure 1: **a.** For each lambda value between 1 and 100, we drew 1,000,000 samples from the corresponding Poisson distribution. We estimated the percentage of samples that are smaller, equal, or larger than the mean of the distribution. On the left, we represented these percentages as a stacked area chart. On the right, we displayed the difference between the percentages of samples smaller than the mean and larger than the mean. There is a clear bias towards lower values due to the asymmetry of the Poisson distribution. This demonstrates how using a single sample as estimator of the mean introduces a bias towards lower values. **b.** Percentiles corresponding to the mean in the Poisson distribution for different values of the mean parameter.

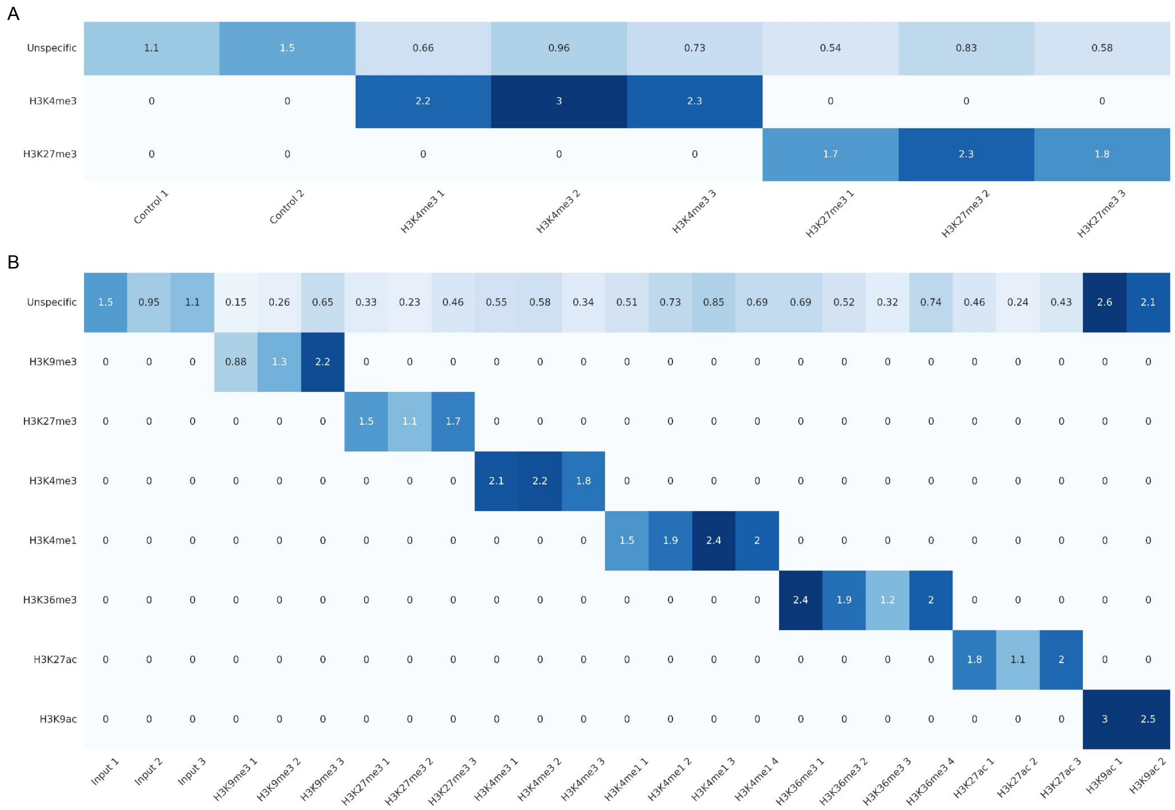


Figure 2: **Mixing matrices from NMF step of DecoDen:** Figures A and B show the mixing matrices for deeply sequenced E114 and E005 respectively. The top row illustrates how much of a cell-type specific signal is contained in each ChIP replicate. All other rows indicate how much histone specific signal contained by a given sample.

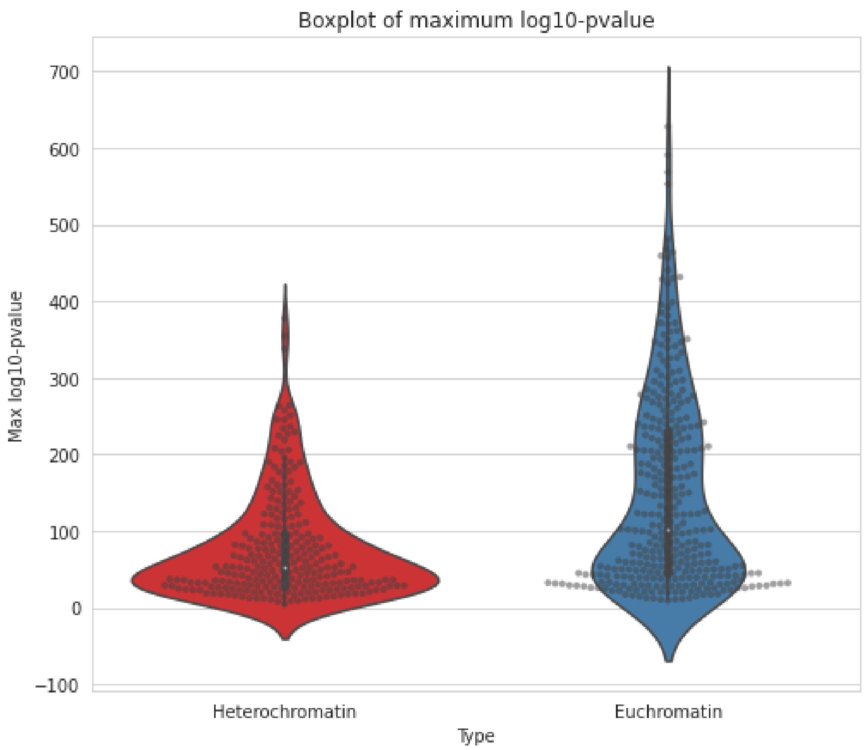


Figure 3: **Violinplot of maximum $-\log_{10} p$ for different tracks in the Roadmap Epigenome dataset and grouped activating and repressive marks separately.** Overall, repressive marks (H3K9me2, H3K9me3 and H3K27me3) have a lower maximum significance compared to activating marks (H3K9ac, H3K4me2, H3K4me3 and H3K36me3). The signal is restricted to chromosome 21 and binned to 25 bp width.

### 3 Supplementary Tables

### 4 Accession Numbers

The accession numbers for the datasets downloaded from the ENCODE portal are as follows - ENCFF200TOF, ENCFF612OXU, ENCFF949VKE, ENCFF571WYJ, ENCFF065HVY, ENCFF061TSZ, ENCFF404DOT, ENCFF779XRN, ENCFF228ABC, ENCFF623DRR, ENCFF346JZT, ENCFF234NNJ, ENCFF124FBL, ENCFF498OUP, ENCFF845XVA, ENCFF644UTO, ENCFF292DMA, ENCFF941XUX, ENCFF696EYT, ENCFF038LZU, ENCFF691LXT, ENCFF602KJS, ENCFF654CCW, ENCFF658CNF, ENCFF992YVV, ENCFF926EQN, ENCFF176HCT, ENCFF526GOB, ENCFF983WQI, ENCFF959PXE, ENCFF090EKJ, ENCFF997VIZ, ENCFF219ZIO, ENCFF612HRT.

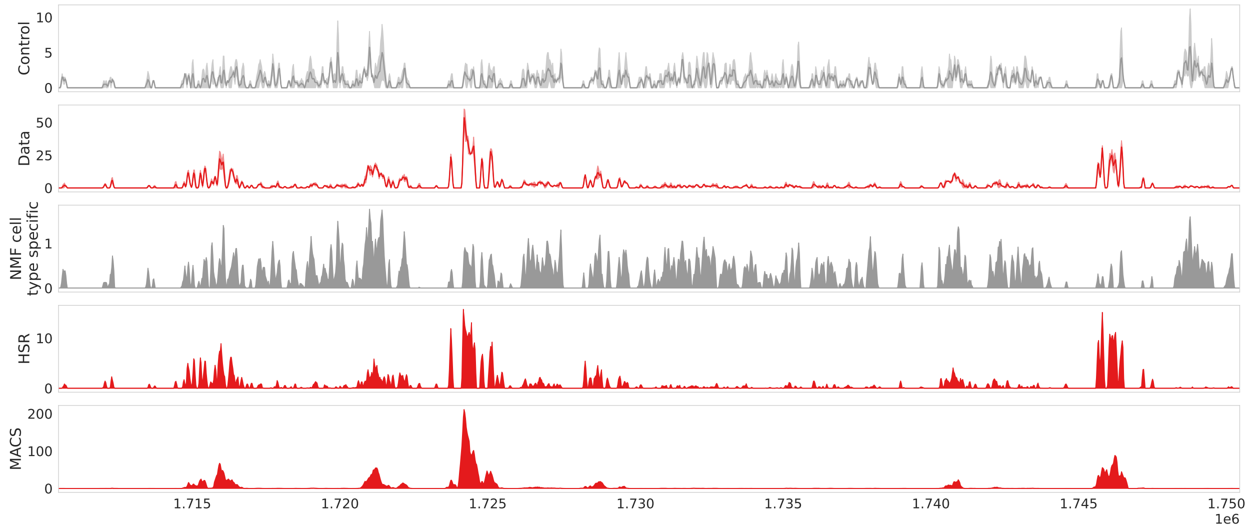


Figure 4: **MACS overestimates the significance of certain regions.** Consider the two peaks at 1.725 MB and at 1.745 MB. Even though the control and treatment coverages look similar, MACS overestimates the significance for the first one (p-value in the order of $10^{-200}$). This distorts the data for downstream tasks. The data used here is for individual 3 (37M) from the ENCODE ENTEx Dataset.

| Histone | Individual | Correlation |
|---|---|---|
| H3K27ac | 37m | 0.90 |
| | 51f | 0.82 |
| | 37m-51f | 0.85 ± 0.04 |
| H3K4me3 | 37m | 0.93 |
| | 51f | 0.87 |
| | 37m-51f | 0.88 ± 0.03 |
| H3K4me1 | 37m | 0.68 |
| | 51f | 0.35 |
| | 37m-51f | 0.47 ± 0.10 |
| H3K27me3 | 37m | 0.50 |
| | 51f | 0.32 |
| | 37m-51f | 0.38 ± 0.05 |
| H3K36me3 | 37m | 0.30 |
| | 51f | 0.27 |
| | 37m-51f | 0.23 ± 0.03 |

Table 1: Table showing correlation between replicates of same individual and between different individuals. This comparison is done for individuals 3 and 4 from the ENTEx Dataset. Note that correlation is calculated only for genomic bins with a coverage greater than a 1. This is done to focus on the foreground, and avoid correlating background signal. Each experimental condition has two technical replicates.
